# Checkpoint activation by Spd1: a competition-based system relying on tandem disordered PCNA binding motifs

**DOI:** 10.1101/2023.05.11.540346

**Authors:** Johan G. Olsen, Andreas Prestel, Noah Kassem, Sebastian S. Broendum, Hossain Mohammad Shamim, Signe Simonsen, Martin Grysbæk, Josefine Mortensen, Louise Lund Rytkjær, Gitte W. Haxholm, Riccardo Marabini, Antony M. Carr, Ramon Crehuet, Olaf Nielsen, Birthe B. Kragelund

## Abstract

DNA regulation, replication and repair are processes fundamental to all known organisms and the sliding clamp proliferating cell nuclear antigen (PCNA) is central to all these processes. S-phase delaying protein 1 (Spd1) from *S. pombe*, an intrinsically disordered protein that causes checkpoint activation by inhibiting the enzyme ribonucleotide reductase, has one of the most divergent PCNA binding motifs known. Using NMR spectroscopy, *in vivo* assays, X-ray crystallography, calorimetry, and Monte Carlo simulations, an additional PCNA binding motif in Spd1, a PIP-box, is revealed. The two tandemly positioned, low affinity sites exchange rapidly on PCNA exploiting the same binding sites. Increasing or decreasing the binding affinity between Spd1 and PCNA through mutations of either motif compromised the ability of Spd1 to cause checkpoint activation in yeast. These results pinpoint a role for PCNA in Spd1-mediated checkpoint activation and suggest that its tandemly positioned short linear motifs create a neatly balanced competition-based system, involving PCNA, Spd1 and the small ribonucleotide reductase subunit, Suc22^R2^. Similar mechanisms may be relevant in other PCNA binding ligands where divergent binding motifs so far have gone under the PIP-box radar.

## Introduction

In Archaea and Eukaryota, the sliding clamp proliferating cell nuclear antigen (PCNA) is found at the centre of all DNA replication, modification and repair processes (1–4). A similar molecule, the β-clamp, has the same basic function in Bacteria (4, 5). These clamps are essential to all living cells. PCNA is a homotrimer in the shape of a ring, the centre of which can harbour a DNA double strand. The trimer is latched onto DNA by a clamp loader in an ATP dependent manner (6). The ring then slides along the DNA double strand, carrying relevant interaction partners such as polymerases, replication factors, and nucleases (7). When tethered to PCNA, they are arranged optimally for interaction with DNA and in particular ways relevant to their function (3, 8).

PCNA ligands that do not interact with DNA also exist. They take on other roles, such as in cell cycle regulation (9, 10). Several cytosolic PCNA binding proteins have been identified, some of which are part of the mitogen activated protein kinase (MAPK) signalling cascade, while others are involved in metabolic regulation ((11) and references therein). Once bound to PCNA, some ligands can be recognized by the Cdt2 protein, which is part of the CRL4^Cdt2^ E3 ubiquitylation complex, whereby they become destined to ubiquitylation, and thus, PCNA binding can promote timely degradation of a subset of its ligands (12).

The large number of interaction partners makes PCNA a generic molecular hub, and its pivotal role to life is underlined by the fact that PCNA molecules from Eukaryota and Archaea have been separated in their respective domains for at least 2.5 billion years, yet their detailed molecular structures have remained astonishingly similar (9). PCNA has one well-documented central ligand binding site in each subunit, constituted by a hydrophobic- and a Q-pocket, which binds a wealth of ligands that differ broadly in sequence. Most ligands are intrinsically disordered in their free state and adopt a single 3_10_- helical turn when bound (6–8, 13, 14). Although very rare, a few exceptions exist, where the ligand either forms an extended strand of a stretch of residues (e.g., Activator 1 40 kDa subunit complex (6)) or is α-helical (e.g., Pif1 (15)), or where ligands also bind to other patches on PCNA (16).

The central PCNA-binding residues in the ligands constitute a short linear motif (SLiM) and are referred to as the *PCNA interacting peptide* (PIP) (17). The PIPs are divided into at least three groups; the PIP-box, the PIP-degron (18, 19) and the AlkB homologue 2 PCNA-interacting motif (APIM) (20). These three groups have the following consensus motifs: PIP-box: QXXψXXΩΩ, PIP-degron: QXXψTDΩΩXXX[+] and APIM: [+]Ωψψ[+], where X is any amino acid, ψ is an aliphatic residue, Ω is an aromatic residue and [+] is either K or R. None of the residues are completely conserved throughout all PCNA binding SLiMs or exclusively necessary for binding, reflecting a high degree of diversity and adaptability, and thus a broad tolerance by PCNA for accommodating different sequence motifs (8).

For reasons that so far have been somewhat elusive, some PCNA ligands have been shown to contain more than one PCNA binding SLiM (21). The transcription factor TFII-I is active in translesion DNA-synthesis through binding to PCNA via four APIM motifs (22). In the case of human DNA polymerase (Pol) η, three PIP-boxes (PIP1-3) were discovered (23). Here, the presence of PIP3 and to a lesser degree PIP2 promoted monoubiquitylation of PCNA in response to DNA damage (23). Pol η polymerase activity was dependent on the presence of PIP1 and PIP3, but not PIP2, so tandemly repeated PIP-boxes may contain specialised SLiMs, the binding of which depend on the cellular state. Whether the three Pol η PIP sites bind in a cooperative manner or bind to different PCNA subunits remains to be investigated.

One of the PCNA ligands unreported to interact with DNA is S-phase delaying protein 1 (Spd1) from *Schizosaccharomyces (S.) pombe*. Spd1 is a small intrinsically disordered protein (IDP) (24, 25), originally identified as an inhibitor of DNA replication (26, 27). Subsequent studies established that Spd1 inhibits the enzyme complex ribonucleotide reductase (RNR) by several complex mechanisms (25), one of which is to sequester the small subunit Suc22^R2^ inside the nucleus. During DNA synthesis, Spd1 becomes degraded by a mechanism depending on the CRL4^Cdt2^ ubiquitylation pathway. Failure to degrade Spd1 renders cell survival reliant on activation of the Rad3^ATR^ checkpoint, presumably because of starvation for dNTP DNA building blocks. The Spd1 protein is highly unstable (28) and the coupling of its degradation with ongoing replication may serve to ensure optimal dNTP levels for error free DNA synthesis via feedback inhibition of RNR (29, 30).

Consistent with its CRL4^Cdt2^ dependent degradation, Spd1 was shown to contain a PCNA-binding SLiM of the PIP-degron type constituting residues Q30-K42 (29), and by bimolecular fluorescence complementation (BiFC), Spd1 was demonstrated to interact with PCNA *in vivo* (31). These studies reported a modest but significant residual binding to PCNA with Spd1^44-124^ (*i.e.*, in the absence of the PIP-degron), suggesting the presence of additional PCNA binding by Spd1. However, the sequence of the Spd1 PIP-degron makes it one of the most divergent among 80 recently curated ligands (8), in that it lacks both aromatic residues of the motif, **Fig. 1A**. One of these is even substituted by a glycine (Gly37). As this region, which encompasses the PIP-degron, directly overlaps with the HUG domain shown to be implicated in sequestering the small RNR subunit Suc22^R2^ into the nucleus (25), this might, in part explain the divergence from the PIP-degron consensus.

**Fig. 1.**
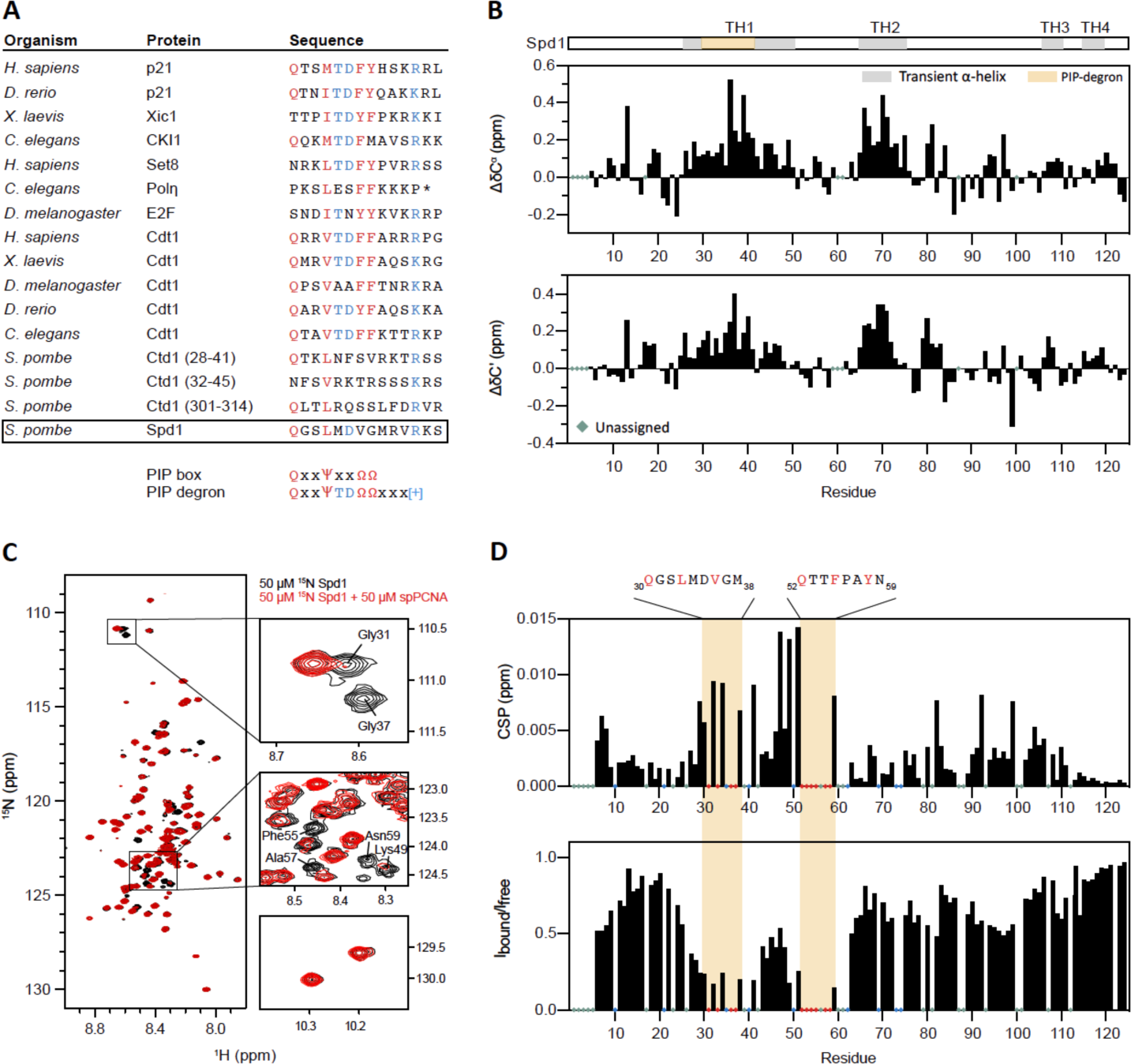
Spd1 carries not one, but two PIP motifs. **A**) Alignment of 16 different PIP-degron SLiMs from Fungi and Animalia illustrates a remarkable conservation. The PIP-degron of Spd1 is permutated almost beyond recognition and lacks two canonical aromatic residues. **B**) Secondary chemical shift of C^α^ (upper panel) and C’ (lower panel) nuclei of Spd1 in solutions calculated from assignment in the absence and presence of 8 M urea. Transient helices are indicated by TH1 through TH4. **C**) ^1^H-^15^N-HSQC spectra of ^15^N-labelled Spd1 in the absence (black) and presence (red) of equimolar concentration of *S. pombe* PCNA. Signals from representative residues that disappear from the spectrum in the presence of PCNA are highlighted to the right. **D**) Chemical shift perturbations (upper panel) and intensity ratios (lower panel) of ^15^N-Spd1 upon addition of PCNA. Two regions are affected, the known PIP-degron N-terminally, and 14 residues C-terminal to that, a hitherto unidentified PIP-box.

In the present paper, we analyse PCNA binding by Spd1 in detail using Spd1, Spd1-derived peptide ligands, as well as ligand variants. We apply nuclear magnetic resonance (NMR) spectroscopy, isothermal titration calorimetry (ITC), *in vivo* assays, X-ray crystallography, and Monte Carlo (MC) simulations analyses. We identify an additional PCNA binding motif in Spd1, a PIP-box, and show that the two low-affinity, tandemly positioned motifs exchange rapidly on PCNA *in vitro.* Computational simulation analyses suggest that in the PIP-degron-bound complex, the PIP-box binds to the interdomain connecting loop on the outer surface of PCNA, whereas in the PIP-box-bound state, the PIP-degron binds to the interior cavity of PCNA. Increasing or decreasing the affinity between Spd1 and PCNA through mutations caused a reduction in the ability of Spd1 accumulation to cause Rad3^ATR^ activation *in vivo*. The results reveal an example of a neatly balanced competition-based system at the heart of checkpoint activation, involving PCNA, Spd1, Suc22^R2^, and most likely also other factors in the DNA replication network. We speculate that this could be relevant for many other PCNA binding ligands, which so far have gone under the PIP-box radar.

## Materials and methods

### Protein purification and peptides

Spd1 and variants were expressed and purified from ^15^N-labeled (^15^NH_4_SO_4_) of ^15^N and ^13^C labelled (^13^C_6_-glucose) M9 medium as previously described (25) and NMR assignments of Spd1 obtained from BMRB code 51376 (32).

PCNA was purified using two different approaches. PCNA was expressed from a modified pET24b vector with His_6_-SUMO-spPCNA after transforming into *E. coli* BL21(DE3) cells and grown on an agar plate containing 50 μg/mL kanamycin (kan) overnight (o/n) at 37 °C. A single colony from the plate was transferred to 10 mL of room temperature (RT) LB medium containing 50 μg/mL kan and incubated at 37 °C o/n while shaking at 180 rpm. The overnight culture was added to 1 L LB medium containing kan to a final concentration of 50 μg/mL and the culture incubated at 37 °C while shaking at 180 rpm. At OD_600_ ∼0.8, isopropyl-d-thio-Galactoside (IPTG) was added to a final concentration of 0.5 mM to induce expression and the culture left o/n at 20 °C shaking at 180 rpm. For NMR experiments, ^2^H, ^15^N, ^13^C labelled PCNA was expressed in deuterated M9 media containing 2 g/l ^13^C, ^2^H labelled glucose and 1 g/l ^15^NH_4_Cl in 98% D_2_O with the addition of kan to a concentration of 25 ug/ml. Overnight cultures in LB medium were made as described above. Precultures were made by adding 500 μl of overnight culture to 20 ml deuterated M9 media with 25 μg/ml kan and the precultures were grown o/n at 37°C shaking at 180 rpm. The preculture was added to 480 ml deuterated M9 media with 25 μg/ml kan and incubated at 37°C, 180 rpm. At At OD_600_ ∼0.5 the cells were induced by adding IPTG to a final concentration of 1 mM and the culture was left left o/n at 20 °C shaking at 180 rpm. The cells were harvested by centrifugation at 5000 x g for 15 min, drained and the pellets stored at −20 °C. The cell pellets were thawed on ice and resuspended in 40 mL of solution A (50 mM Tris-HCl pH 8.0, 150 mM NaCl, 10 mM imidazole, 10 mM β-mercaptoethanol (bME)) supplemented with a cOmplete^™^, EDTA-free Protease Inhibitor Cocktail (Roche). The sample was applied to a French press cell (40 ml), prechilled to 4 °C and the cells were pressurized to approximately 20000 psi. The cell lysate was collected on ice. Cell debris was removed by centrifugation at 20000 x g for 30 min at 4 °C. The supernatant was added to a 50 mL Falcon tube containing 5 ml Ni-NTA suspension (Qiagen) resuspended in solution A and left to settle on a tilting table for 1 hour at RT. The solution was loaded onto a disposable plastic chromatography column prior to being washed with 50 mL solution B (1 M NaCl, 50 mM Tris pH 8.0, 10 mM imidazole, 10 mM bME). The column was washed with 50 mL ice-cold solution A,and eluted using 10 mL of solution C (250 mM imidazole, 150 mM NaCl, 50 mM Tris pH 8.0, 10 mM bME). The the imidazole was removed by exchanging the buffer to solution D (50 mM Tris, 150 mM NaCl, 10 mM bME, pH 8) using an Amicon 10.000 MWCO 15 ml spin filter and ∼0.4 mg of ULP1 protease was added to cleave off the His-SUMO tag at room temperature o/n. Cleaved PCNA was separated from His-SUMO and uncleaved His-SUMO-PCNA by incubating the protein solution with 5 ml Ni-NTA suspension (Qiagen) resuspended in solution D on a tilting table for 30 min. The solution was loaded onto a disposable plastic chromatography column and the flowthrough collected. The flowthrough was concentrated to ∼3 mL using an Amicon 10.000 MWCO 15 ml spin filter. The samples were centrifuged at 20.000 g for 15 min. to remove precipitated protein before being applied to a size exclusion chromatography column (HiLoad 16/60 Superdex 200, 120 mL) equilibrated with 2 CV of 20 mM Sodium phosphate, 100 mM NaCl, 1mM DTT, pH 7.4. The protein was purified at a flowrate of 1.0 mL/min and eluted over 1.5 CV. The column was run on an ÄKTA FPLC apparatus and all buffers were sterile filtered (0.22 μM filter) before use. After analysis by SDS-PAGE, relevant fractions were pooled and stored at 4 °C.

To express unlabelled PCNA, M15 [pREP4] *E. coli* cells from a glycerol stock stored at −80◦C, containing the PQE32 vector with the coding region for *S. pombe* PCNA inserted, were grown on an agar plate containing 100 μg/mL ampicillin (amp) and 50 μg/mL kan o/n at 37 °C. The expressed PCNA contained an N-terminal His_6_-tag as well as cloning remnants and thus had the N-terminal sequence of MRGSHHHHHHGIP, i.e., 13 additional N-terminal residues. A single colony from the plate was transferred to 10 mL of room temperature (RT) LB medium containing 100 μg/ml amp and 100 μg/mL kan and incubated at 37 °C o/n while shaking at 180 rpm. The overnight culture was added to 1 L LB medium containing amp and kan to final concentrations of 100 μg/mL and 50 μg/mL, respectively and the culture incubated at 37 °C while shaking at 180 rpm. At OD_600_ 0.7, IPTG was added to a final concentration of 1 mM to induce expression and the culture left for five hours at 37 °C shaking at 180 rpm. The cells were harvested by centrifugation at 4000 x g for 20 min, drained and the pellets stored at −20 °C. The cell pellets were thawed on ice and resuspended in 40 mL of solution A (50 mM Tris-HCl pH 8.0, 150 mM NaCl, 20 mM imidazole). The sample was applied to a French press cell (40 ml), prechilled to 4 °C. The cells were pressurized to approximately 10000 psi and the outlet flow rate adjusted to 1 drop/s. The cell lysate was collected on ice. Cell debris was removed by centrifugation at 30000 x g for 20 min at 4◦C. The supernatant was added to a 50 mL Falcon tube containing 5 ml Ni-NTA suspension (Qiagen) resuspended in 2 mL solution A and left to settle on a tilting table for 1 hour at RT. The solution was loaded onto a disposable plastic chromatography column prior to being washed with 120 mL solution B (1 M NaCl, 50 mM Tris pH 8.0, 20 mM imidazole). The column was washed with 80 mL ice-cold solution A, eluted using 10 mL of solution C (250 mM imidazole, 50 mM Tris pH 8.0), and concentrated using a 10000 kDa molecular weight cut-off (MWCO) centrifugal filter (Millipore) until the volume was 1.5 ml. It was applied to a size exclusion chromatography column (Sephacryl S-200 HR 16/60, 120 mL) equilibrated with 2 CV of solution D (10 mM NaH_2_PO_4_, pH 7.4, 100 mM NaCl, 2 mM DTT) at a flowrate of 1.0 mL/min and eluted over 1.5 CV. The column was run on an ÄKTA FPLC apparatus and all buffers were sterile filtered (0.22 μM filter) before use. After analysis by SDS-PAGE, relevant fractions were pooled, flash cooled using liquid nitrogen and stored in the freezer at −20◦C until further use.

Peptides covering the PIP-box (27-46) and the PIP degron (49-68) of Spd1 were bought N-terminally acetylated and C-terminally amidated from Synpetide Co Ltd (Shanghai, China) and KJ ROSS-PETERSEN ApS (Copenhagen, Denmark), respectively. The purity was > 95% according to HPLC and mass spectrometry spectra recorded of both peptides.

### NMR spectroscopy

All NMR samples were prepared by adding 10 % (v/v) D_2_O, 0.02 % (w/v) sodium azide and 0.5 mM DSS to 309 μL of a protein solution in 10 mM NaH_2_PO_4_ and 100 mM NaCl (pH 7.4) in the presence and absence of ∼8 M urea. The pH of the samples was checked and regulated to 7.4 with appropriate concentrations of hydrochloric acid and sodium hydroxide if needed. To remove insoluble material, the sample was centrifuged for 5 min at 20000xg. The samples were transferred to 5 mm Shigemi NMR tubes.

All NMR spectra were recorded on a Varian Unity Inova 750 or 800 ^1^H MHz NMR spectrometers or on a Bruker Avance III HD 600 MHz spectrometer with a Bruker proton-optimized quadruple resonance NMR ‘inverse’ (QCI) cryoprobe. Pulse sequences used were from the Varian Biopack or Bruker Topspin. ^1^H chemical shifts were referenced to DSS and the ^15^N and ^13^C chemical shifts indirectly using the gyromagnetic ratios. All spectra were zero-filled, apodized using cosine bells in all dimensions, Fourier transformed, and the zero-order phase was corrected manually using either nmrDraw, a component of NmrPipe (33) or qMDD if spectra were recorded using non-linear-sampling (NLS). All spectra were analysed using the Analysis software. ^1^H-^15^N-HSQC spectra and triple resonance spectra, HNCACB CBCACONH (*99*), HNCO, HNCACO and HNN were recorded for Spd1 at 4° C in the presence and absence of 7M urea. *S. pombe* PCNA was assigned from a ^2^H-^13^C-^15^N labelled sample at monomer concentration of 1 mM using the deuterium decoupled TROSY variants of HNCA, HNCO, HNCACB, HNcoCA, HNcaCO as well as a 15N edited 3D-TROSY-NOESY to correlate spatially close amide protons. All assignment experiment were recorded on a 750 MHz Bruker Avance III HD Spectrometer at 35 °C. The assignment was deposited to the BMRB with the deposition number 51437. All backbone experiments were recorded using NLS with a data reduction to 20 %.

### Secondary chemical shifts (SCSs) and chemical shift perturbations (CSPs)

To calculate the secondary chemical shifts for Spd1 the following equation was used

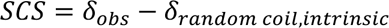

where the chemical shifts of Spd1 in 8 M urea was taken as the intrinsic random coil chemical shifts. To quantify the CSPs of the signals from addition of binding partners, ^1^H,^15^N-HSQC spectra were recorded at increasing concentrations of added ligand and the normalized CSPs from individual peaks calculated using as the weighted Euclidean distance

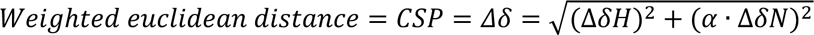

Where α is 0.154 (34).

### Determination of *K_d_*s from NMR titrations

^1^H-^15^N-TROSY-HSQCs of 150 μM ^15^N-^13^C-^2^H spPCNA were measured a Varian Unity Inova 800 MHz Spectrometer with increasing concentrations of the Spd1^27-46^ and Spd1^49-68^ peptides (up to 1.9mM and 2.2 mM respectively). The CSPs were extracted from the CCPN Analysis software program used for spectra analysis (35). and plotted against the ligand concentration for 5 non overlapping signals from residues that showed the largest CSP. The data was fitted assuming a two-state binding model with the formula:

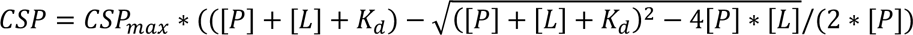

where *CSP_max_* is the chemical shift perturbation upon saturation and [P] and [L] are the PCNA and peptide concentration respectively.

### Diffusion measurements by NMR to obtain *R_h_*

^15^N-edited 1D ^1^H NMR spectra of 50 μM ^15^N-labelled Spd1 were recorded at 25°C on a Bruker 600 MHz spectrometer equipped with a QCI-cryoprobe and Z-field gradient with varying gradient strengths from 2.4 to 47.2 G/cm, a diffusion time Δ of 100 ms and gradient length δ of 4 ms. Translational diffusion coefficients D for Spd1 (50μM) were determined by fitting the peak intensity decay of the amide resonances in Dynamics Center v2.5.6 (Bruker, Switzerland) using the Stejskal-Tanner equation (36). R_h_ was calculated from the diffusion coefficient using the Stokes–Einstein relation R_h_ = k_B_T/(6πηD) with η = 0.890 mPa * s (dynamic viscosity of water at 25°C).

### Crystallization and data collection

PCNA was concentrated to 0.33 mM (10 mg/ml) (based on absorption at 280 nm and a theoretical extinction coefficient) in buffer A (100 mM NaCl, 10 mM NaH_2_PO_4_ pH 7.4, 2 mM DTT). Lyophilized Spd1^27-46^ and Spd1^53-64^ (N-terminally acetylated and C-terminally deamidated) purchased from Synpeptide Co Ltd. were solubilized in buffer A to reach a concentration of 3.8 mM. The PCNA and Spd1^27-46^ or Spd1^53-64^ solutions were mixed 1:1 (by volume) prior to crystallization. Hanging drop vapor diffusion experiments were set up manually in Hampton Research VDXm plates containing 24 wells with siliconized circular glass cover slides. To crystallize PCNA in complex with either the PIP-degron, (Spd1^27-46^) or the PIP-box (Spd1^53-64^), 2 μL of the mixture (0.17 mM PCNA and 1.9 mM Spd1^27-46^) was mixed with 2 μL reservoir solution containing 100 mM TRIS-HCl pH 7.5, 18% (w/v) PEG4000, 6% (v/v) glycerol. All stock solutions were filtered through a 0.22 μm filter, except for (PEG) 4000 due to the high viscosity. Crystals were left to grow for 24 hours, then harvested and flash-cooled in liquid nitrogen. Data were collected at the European Synchrotron Radiation Facility (ESRF), beamline ID23-1, equipped with a Pilatus detector (3M, S/N 24-0118). Each frame (Δφ=0.1°) was exposed for 0.2 s with a monochromatic beam, λ=0.8729 Å. Obtaining data to a sufficient resolution (better than 3 Å) proved challenging. The diffraction power often varied depending on which part of a crystal was exposed to the X-ray beam, and there were radiation damage problems. Prior to collecting an entire dataset, various parts of each crystal were tested, and the strategy program EDNA (37) was run on four images, thus assessing radiation damage and estimated lifetime of the crystal.

### Structure solution

Phases for the structure factor amplitudes were calculated using molecular replacement. The search model was the crystal structure of trimeric human PCNA (51.3 % sequence identity, PDB accession code 3WGW). The search was performed using Phaser (all programs mentioned are part of the CCP4i suite(38) unless otherwise stated) and resulted in one solution. Molecular replacement was followed by a rigid body refinement in Refmac5 (39). The amino acid sequence was altered manually in the macromolecular graphics program Coot (40). Several loop residues were ill (or not at all) defined in the electron density. These regions were omitted from the model and the majority if the initially missing residues were gradually built back into the model in iterative rounds of building and phasing, as the electron density map improved. Non-crystallographic symmetry constraints were imposed between the three subunits early in the building/refinement procedure, greatly improving the phases. Each refinement round was performed using jelly-body refinement with sigma set to 0.005 (tight restraints) and non-crystallographic symmetry restraints, using the ‘tight main chain & medium side chain’ restraints option. The final R_work_/R_free_ values were 0.23 and 0.26, respectively, to 2.9 Å resolution. Some residues were omitted from the model because of missing or noisy electron density, especially in the interdomain connecting loop (IDCL) (residues 120-130) and a loop around residues 185-193. Statistics and crystallographic details are given in Table S1. The model and structure factors (electron density) are deposited with the Protein Data Bank, accession code 6QH1.

### Molecular dynamics simulations

We performed Monte Carlo simulations to visualize the conformations of Spd1 in the PCNA bound state. We used the ABSINTH implicit solvent model (41) as implemented in the Campari software (42). We simulated two different systems, in which Spd1 was either bound to PCNA *via* the PIP-degron or the PIP-box motif. Residues, which were missing in the crystal structure (PDB accession code 6QH1), were added and a starting structure was generated, both using Modeller (43). In the simulations, the PCNA structure is fixed except for the eight C-terminal residues which belong to a disordered region. The six residues interacting directly with the generic ligand binding site on PCNA were also fixed. These residues are GSLMDV (the PIP-degron site) and TTFPAY (the PIP-box). A harmonic potential was chosen to keep the Spd1 C^α^ atoms of the 3 residues in the binding site closest to PCNA restrained, to avoid unbinding of Spd1. We added ions to neutralize the system and simulated two different NaCl concentrations, 5 mM and 20 mM. These numbers are lower than experimental values because higher salt concentrations rendered the Monte Carlo sampling computationally too slow. We simulated both systems with Replica Exchange (44), using 36 replicas spanning the range from 300K to 400K. The analysis of the trajectories was done with Matplotlib (45) and MDtraj (46).

### Isothermal titration calorimetry

Purified PCNA was buffer exchanged into an ITC buffer (10 mM sodium phosphate, 100 mM NaCl, 1 mM TCEP, pH 7.4) and concentrated using an Amicon 10.000 MWCO 15 ml spin filter. The lyophilized 2F2 peptide was solubilized in ITC buffer and the pH adjusted to 7.4. The ITC experiment was repeated three times with spPCNA in the cell and the 2F2 peptide in the syringe. The concentration of the monomeric spPCNA ranged between 23.2 µM and 25.9 µM, and the 2F2 peptide concentration was between 243 µM and 264 µM as determined on a NanoDrop ND-1000 spectrophotometer using λ = 280 nm absorbance. Protein and peptide samples were centrifuged at 20.000 x g for 15 min. at 37 °C to degas the samples. Data were recorded at 37 °C on a Malvern MicroCal PEAC-ITC instrument with a stir speed of 750 rpm. Data were analysed using the MicroCal PEAQ-ITC Software and fitted with a one site binding model.

### Yeast molecular genetics and cell biology

The *S. pombe* strains used in the present study are listed in Supplementary **Table S2**. Spd1 mutants were generated by site-directed mutagenesis, and used to replace *Δspd1*::*ura4^+^* in the genome by homologous recombination, selecting for FOA resistance. Standard genetic procedures were performed according to (47). Physiological experiments, Bimolecular Fluorescence Complementation (BiFC) and spot test survival assays were as described previously (31). GFP-Suc22 nuclear localization was performed as in(24). Fluorescence and Dark field microscopy were performed on a Zeiss Axio Imager platform (Zeiss, Jena, Germany).

## Results

To address the suggested interaction between PCNA and Spd1 at the molecular level and investigate if and how SLiM divergence still enables activation of the checkpoint, we set out to establish the molecular properties and structural propensities of Spd1, the Spd1-PCNA complex, and complexes between PCNA and two Spd1 derived peptides, respectively.

### Spd1 is intrinsically disordered but compact

To enable identification of the exact binding regions between PCNA and Spd1, we first addressed the conformational properties of Spd1. Using NMR spectroscopy, we analysed the chemical shifts of Spd1 at neutral pH in the absence and presence of 7M urea and extracted the secondary chemical shifts (SCS) (**Fig. 1B**). These can reveal stretches adopting transient secondary structure configurations. Positive consecutive SCSs for both C^α^ and C’, indicative of transient helicity (TH), were observed in two major regions TH1 (L25-S50) and TH2 (T65-T75), and two minors ones, TH3 (E106-L110) and TH4 (V115-G119). TH2, −3 and −4 located to regions of Spd1 with so far unassigned function. TH1, however, included the unusual PIP-degron motif in Spd1 (Q30-K42). This distributed content of transient structure made us address whether the Spd1 ensemble tended towards being extended or compact. Using pulsed field gradient NMR diffusion experiments, the radius of hydration, *R*_h_, was extracted. Compared to published scaling laws (Table 1) (48, 49), the hydrodynamic ration, *R_h_* of Spd1 was 27.8 ± 0.2 Å indicating it to be slightly more compact under these conditions than most IDPs (49). The protein also precipitated at concentrations higher than 50 μM, worsened by increasing salt concentration suggesting the presence of hydrophobic interactions formed by both intra and intermolecular contacts.

**Table 1.** Chain dimensions of Spd1.

| | $R_h$ (Å) | Scaling | reference |
| --- | --- | --- | --- |
| NMR experimental | $27.8 \pm 0.2$ | | This work |
| Chemically unfolded | 32.9 | $2.33 * N^{0.549}$ | (65) |
| IDP | 28.96 | $2.49 * N^{0.509}$ | (49) |
| IDP (improved) | 28.61 | $F * 2.49 * N^{0.509}$ | (49) |

N: number of residues. F: factor calculated from the empirical constants A-D and the fraction of prolines and charge excess.

### Spd1 contains two functional PCNA binding sites

Next, we investigated the Spd1:PCNA complex. To identify the PCNA interaction site in Spd1, ^1^H-^15^N-HSQC spectra of ^15^N-labelled Spd1 in the presence and absence of *S. pombe* PCNA were recorded and compared (**Fig. 1C**, Supplementary **Fig. S1**). A large decrease in the NMR peak intensity ratios between bound and unbound Spd1 was observed in the regions covering I29-A71 (**Fig. 1D**). Sets of peaks from two distinct regions fully disappeared upon addition of PCNA, indicating these regions interact with PCNA. One (_30_<u>Q</u>GS<u>L</u>MD<u>VG</u>M_38_), covering the previously suggested PIP-degron and, additionally, a second motif was discovered (_52_<u>Q</u>TT<u>F</u>PA<u>YN</u>_59_), 14 residues downstream of the last residue of the degron. This binding site motif was identified as belonging to the family of PIP-boxes.

Apart from the two PIP SLiMs, other parts of Spd1 showed signs of interacting with PCNA. The C-terminal part (L72 - L124) exhibited only a modest decrease in intensity ratio, but enough to suggest that the dynamics of these residues was affected by the presence of PCNA, although to a lesser degree. Furthermore, the dynamics of a few residues near the N-terminus of Spd1 (R6-M8) also changed, hinting that they too partook dynamically in the interaction. These results showed that complex formation with PCNA caused large changes in the dynamics of Spd1.

### Non-canonical binding of Spd1 revealed by the crystal structure of the PIP-degron-PCNA complex

With the identification of an additional PCNA SLiM in Spd1 with a much stronger resemblance to a canonical PIP-box, the question arose how these two sites target the PIP-binding site on PCNA. We therefore set up crystals of PCNA in complex with two Spd1-peptides containing either the PIP-degron (Spd1^27-46^), the PIP-box (Spd1^49-68^), or full-length Spd1. Whereas no crystals were seen with intact Spd1, crystals grew in the presence of both peptides within 24 hours. Unfortunately, the diffraction data from crystals grown in the presence of Spd1^49-68^ (the PIP-box) did not provide sufficient electron density from the ligand to allow determination of its bound structure.

The structure of the complex of PCNA with Spd1^27-46^ (the PIP-degron) was solved to 2.9 Å resolution with *R_work_* and *R_free_* values of 0.23 and 0.26, respectively (PDB code 6QH1, Supplemental **Table S1**). The asymmetric unit contained one PCNA trimer and one Spd1^27-46^ peptide. PCNA formed the well-known ring-shaped homotrimer (**Fig. 2A**). The outer diameter was ∼85 Å, the inner was ∼32 Å. Each subunit consisted of two topologically identical domains, covalently connected through the interdomain connecting loop. The outer part of the ring or the convex surface was a continuous β-sheet, connecting the two domains within each subunit, as well as the subunits to each other. The inner rim was lined with 12 antiparallel α-helices sprinkled with lysine and arginine residues.

**Fig. 2.**
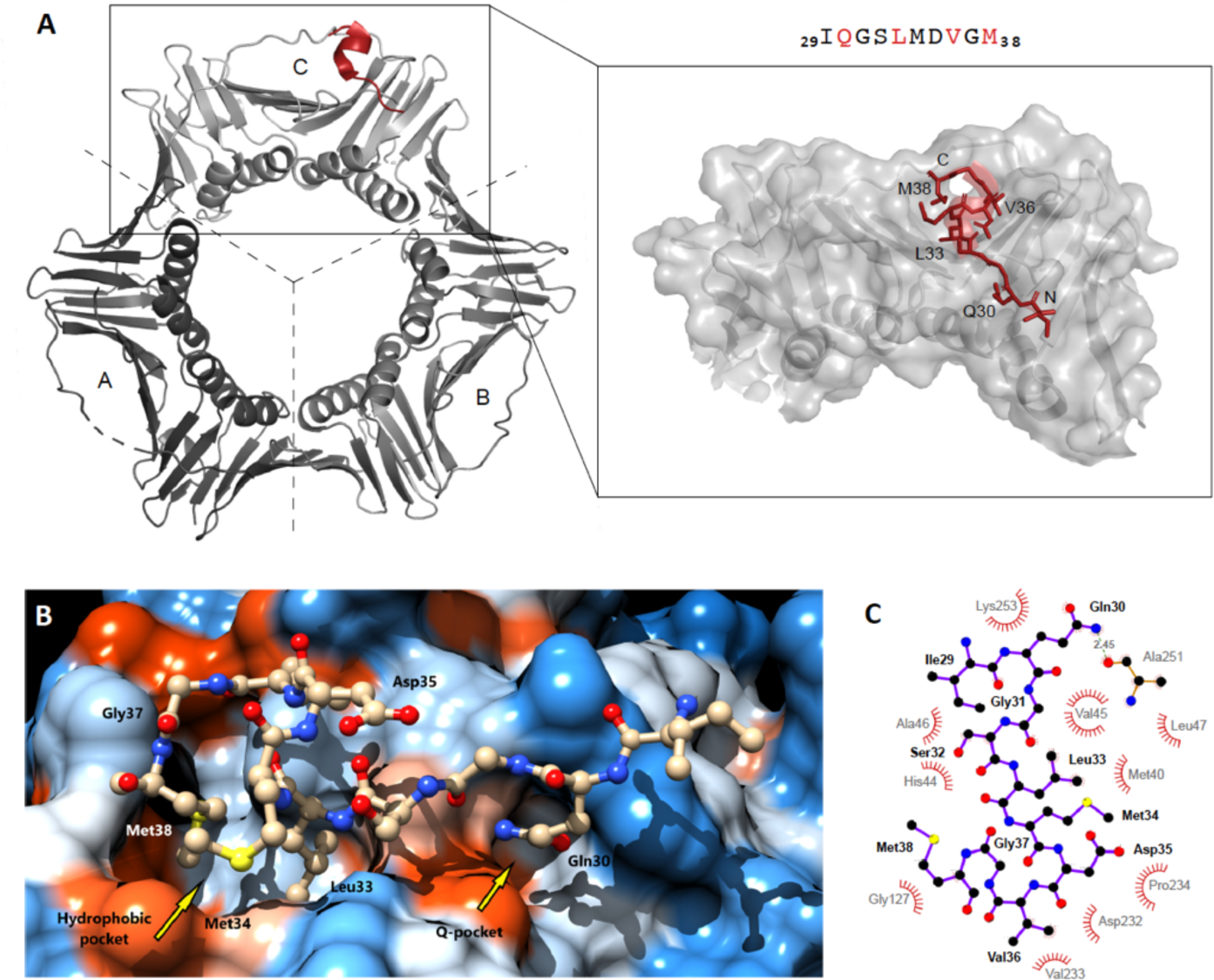
The structure of the PCNA-Spd1^27-46^ complex reveals non-canonical binding. **A**) Ribbon diagram of the homotrimeric PCNA sliding clamp showing the position of the PIP-degron peptide (red). The peptide is extended except for residues Leu33 to Val36, which forms a 3_10_-helical turn in the bound state. **B**) Close up of the Spd1 peptide in the PIP binding pockets. The conserved Gln sits in the Q-pocket on the right, where it makes the only intermolecular hydrogen bond in the complex. The hydrophobic pocket to the left harbours Leu33, Val36 and Met38. **C**) LigPlot (50) representation of the interactions between the peptide and PCNA, highlighting that, apart from the hydrogen bond in the Q-pocket, all interactions are mediated by van der Waals’ contacts.

In the following, the three subunits of the homotrimeric PCNA molecule are referred to as chains A, B, and C, respectively. Of the three ligand binding sites in the crystallised trimer, one (in the B chain) was obscured by a symmetry related PCNA molecule, sterically hindering binding as also observed for other PCNA complexes in the solid state (51, 52). The binding site on the A chain contained residual electron density but not enough to justify modelling. The weakness of the signal presumably owes to very low occupancy of the ligand in the A subunit. However, the ligand could be readily modelled and refined in the remaining binding site on the C chain. Here, the PIP-degron peptide bound in a similar manner to most other published PCNA binding SLiMs, but with a peculiar exception.

Three canonical residues were recognised in the Spd1 PIP-degron motif: the conserved glutamine (Gln30), Leu33 and the degron fingerprint, Asp35. The latter protruded from the 3_10_-helix into the solution as expected (**Fig. 2B**). The canonical glutamine, Gln30, laid, as it does in all cases, in the Q-pocket, where it partook in the only direct intermolecular hydrogen bond in the structure (Gln30 N^ε^ →Ala251 O). The rest of the interactions were van der Waals’ contacts (**Fig. 2C**). The canonical Leu33 was buried in the hydrophobic pockets along with Val36 and Met38. The hydrophobic pockets usually harbour an aliphatic and two aromatic side chains (QXXψTDΩΩXXX). The latter two appear six and seven residues C-terminal to the canonical glutamine. In the present case, however, that part of the pocket was occupied by a methionine (Met38) positioned *eight* residues after Gln30. The eight-rather than seven-residue spacing owes to the insertion of Gly37, which ensures the positioning of Val36 and Met38 where the canonical aromatic residues are normally found. So, the binding interface is very similar to those of canonical binding motifs despite the deviation from the canonical degron signature (53). To our knowledge, this is unique to Spd1.

### The two PCNA-binding motifs of Spd1 bind to and exchange rapidly in the same site

To test whether both motifs in the form of Spd1-derived peptides could bind to PCNA in solution, we expressed ^15^N,^13^C,^2^H-labelled PCNA from *S. pombe* and assigned the backbone chemical shifts (**Fig. 3A**, Supplemental **Fig. S2**). Subsequently, peptides containing the two SLiMs (Spd1^27-46^ and Spd1^49-68^) were analysed one by one for their ability to bind PCNA in solution. This was done by following the chemical shift perturbations of PCNA as a function of peptide concentration. Addition of either peptide resulted in the same residues exhibiting chemical shift changes. This included residues around Ala46, His125, and Ser150 and Leu250 (**Fig. 3B**), consistent with the residues making up the binding pocket (cf. **Fig. 2B** and **C**). Although each peptide on its own induced changes in chemical shifts, these were sometimes in the same direction and sometimes in opposite directions (**Fig. 3C**) reflecting the different chemistry of the peptides. No peaks from PCNA originating from residues outside the binding region or the C-terminus were affected by peptide binding. Thus, the two peptides representing the previously described PIP-degron and the newly identified PIP-box, respectively, both bound to the known PCNA ligand-binding pockets (**Fig. 3C, D**). Next, we repeated the titration experiments with full-length Spd1 (**Fig. 3B**). Although limited by the low solubility, the chemical shifts represented almost an average between the shifts seen when the peptides were added separately, and no additional changes were observed. This suggested that the tandemly positioned motifs interact independently with PCNA, and that they interchange rapidly on a fast NMR timescale.

**Fig. 3.**
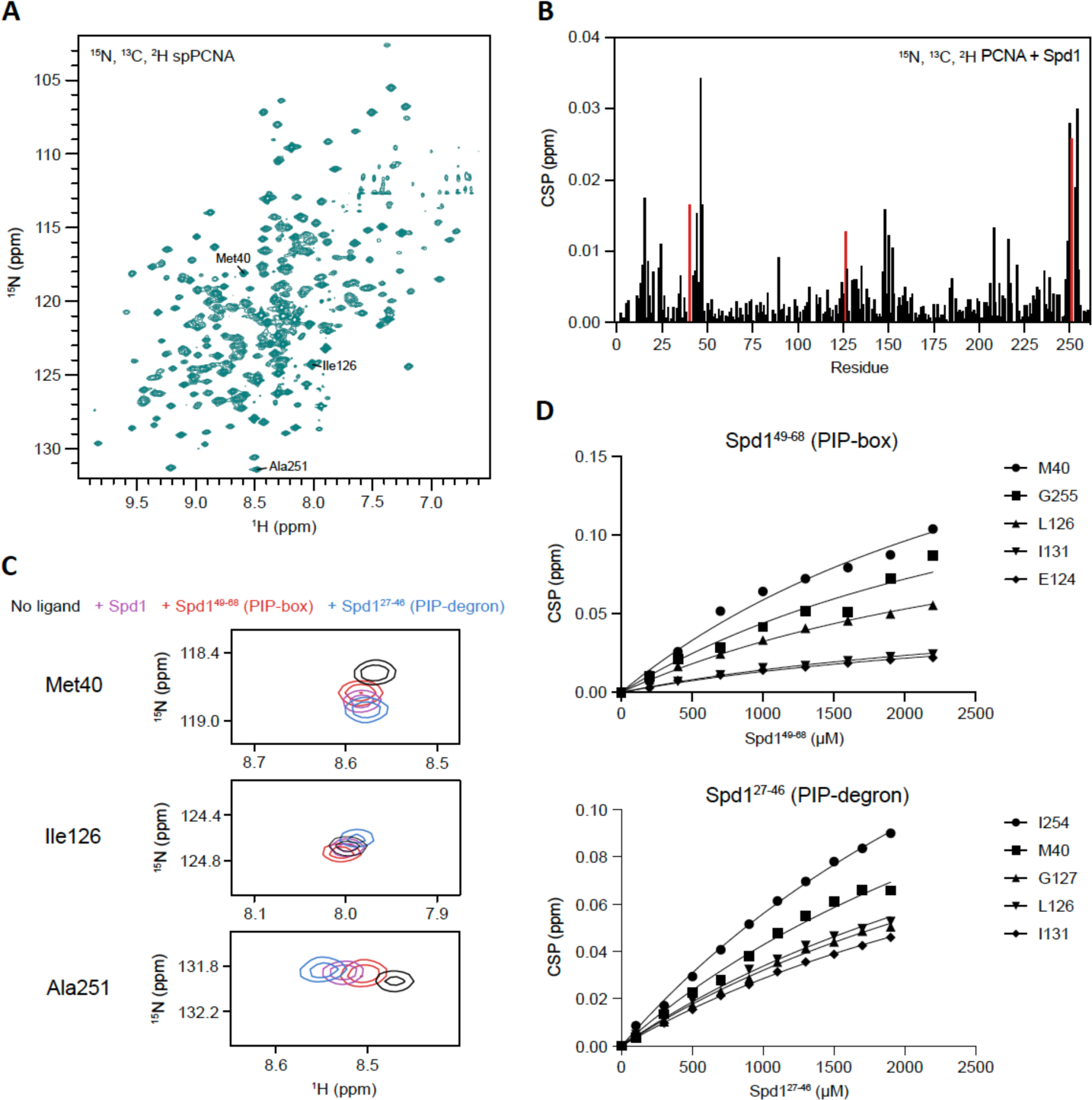
The PIP-degron and PIP-box explore the same binding site on PCNA. **A**) Assigned ^1^H,^15^N-HSQC spectrum of ^15^N,^13^C,^2^H-labeled *S. pombe* PCNA (see also **Fig. S2**). **B**) Chemical shift perturbation of PCNA by the addition of full-length Spd1 (50 μM). The changes in peak position for residues with red bars are shown in detail in C). **C**) Chemical shift perturbations of three representative residues from the three most affected regions of PCNA in the presence of either of the two peptides, Spd1^27-46^ (blue) or Spd1^49-68^ (red) or full-length Spd1 (purple). **D**) Determination of binding affinity, *K_d_*, from changes in chemical shifts for the Spd1-peptides, Spd1^49-68^ including the PIP-box (upper panels) and Spd1^27-46^ including the PIP-degron (lower panels).

The affinity of full-length Spd1 and of the individual peptides for PCNA were then measured by NMR spectroscopy under the assumption that the proximity of the PCNA binding motifs on Spd1 would result in an avidity effect. The affinities for the individual peptides were low, with *K_d_*s of 4.0 ± 0.7 mM and 3.4 ± 1.1 mM for Spd1^27-46^ and Spd1^49-68^, respectively (**Fig 3D**). The magnitude of the chemical shift perturbations using full-length Spd1 were similar to those of the peptides, but at a 10-fold lower concentration. Since a full titration to saturation was not possible because of low Spd1 solubility, the best we could do was to estimate the *K_d_* by comparison of the chemical shifts. By this estimate, the *K_d_* has a value around 300 µM (**Fig. 3D**). Thus, a local concentration effect (54) caused by the proximity of the two PIP SLiMs, contributed to the decrease in the apparent *K_d_* by an order of magnitude rather than a factor of two.

### Implications of binding deficient mutants on checkpoint activation

To investigate the role of the individual PIP sites *in vivo*, we initially made three Spd1 variants. One had two hydrophobic- to glycine substitutions in the PIP-degron (L33G, V36G; *spd1-2G1*), the other two hydrophobic- to glycine substitutions in the PIP-box (F55G, Y56G; *spd1-2G2*), and the last was a combination of the two (*spd1-4G*), **Fig. 4A**. We first repeated the NMR titrations using these three variants and wild-type Spd1 (**Fig. 4C**). The magnitude of the NMR signals from residues in the mutated binding sites did no longer change upon the addition of PCNA, confirming binding deficiency. As expected, no binding to PCNA could be detected for the quadruple variant (**Fig. 4C**). Thus, the NMR data suggested that the two sites are individually capable of binding to PCNA irrespective of the presence of the other and when neither site is present, no binding to PCNA can be detected. We next inserted these mutations into the authentic *spd1* genomic context and tested their ability to cause a requirement for the Rad3^ATR^ checkpoint pathway. When degradation of wild-type Spd1 was repressed by adding thiamine to the *cdt2-TR* allele, cell survival was compromised by the absence of Rad3^ATR^ function at high temperature, as previously reported (31) (**Fig. 4B**). Strikingly. the *spd1-2G2* mutant partially and the *spd1-2G1* mutant fully suppressed the thiamine-induced cell death already at 33°C (**Fig. 4B**). These observations suggested that both motifs in Spd1 contribute to Spd1 mediated checkpoint activation, with the PIP-degron being essential and the PIP-box being important for the ability of Spd1 to cause checkpoint activation.

**Fig. 4.**
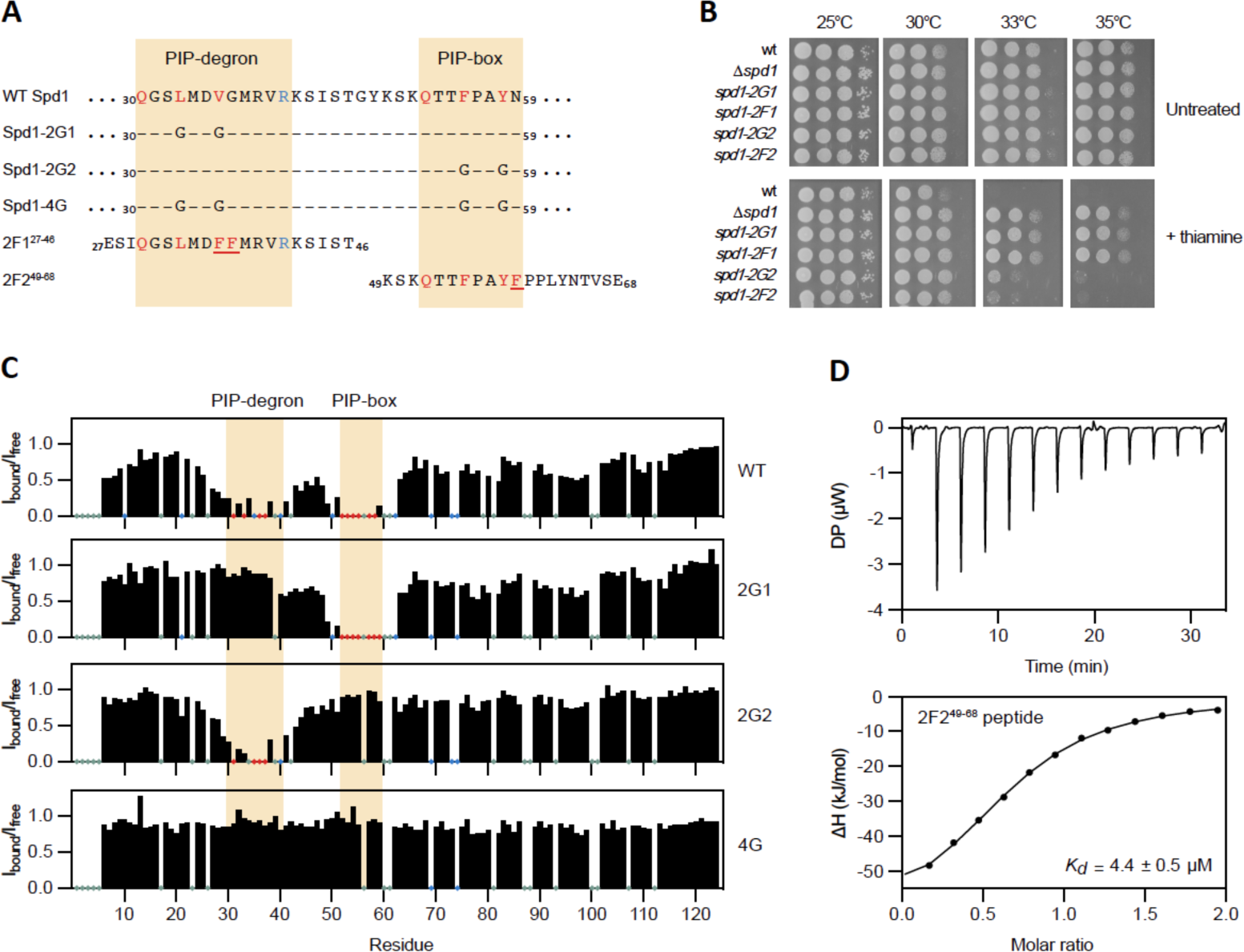
The PIP-box and the PIP-degron of Spd1 act individually and with balanced affinities. **A**) Overview of Spd1 variants. **B**) Serial dilutions of strains with the indicated genotypes spotted on plates either untreated or treated with thiamine and incubated at the indicated temperature for three days. All strains also harbour the *cdt2-*TR and the *rad3-TS* alleles. **C**) NMR peak intensity plots of Spd1 WT, Spd1-2G1, Spd1-2G2 and Spd1-4G from titrations with PCNA. Red dots on x-axis indicate disappearing peaks. **D**) ITC analysis of PCNA binding to the 2F2^49-68^ peptide.

Since Spd1 accumulation causes a requirement for Rad3^ATR^ even when dNTP levels are ensured by an artificial salvage pathway (31), we previously proposed that by binding PCNA, it may block replication by a mechanism similar to the one reported for the p21 protein (31). In the next step we therefore asked if we could restore checkpoint activation by increasing the affinity of the PCNA-Spd1 interaction by introducing canonicity in the motifs. Two variants were constructed in which the two canonical phenylalanine residues of the PIP-box (Y58F;N59F) and PIP-degron (G37F;M38F), respectively were introduced, giving Spd1-2F2 and Spd1-2F1, **Fig. 4A**. Only one of the variants, Spd1-2F2, was soluble *in vitro*, hinting to one reason for the lack of aromatic residues in the PIP-degron. NMR experiments titrating ^2^H,^13^C,^15^N-PCNA with the corresponding 2F2-peptide, showed much larger chemical shift changes in PCNA originating from the ring current effect but also severe line broadening due to intermediate exchange on the NMR time scale, implying a higher affinity. As we could not follow the peaks for a meaningful affinity quantification, the affinity for the 2F2-peptide was instead determined by isothermal titration calorimetry (ITC). This confirmed the much stronger binding to PCNA with a ∼70-fold increase in affinity reaching a *K_d_* of 4.4 ± 0.5 μM (**Fig. 4D**). Surprisingly, when the variants were expressed from the *spd1* locus in fission yeast, a comparable phenotype to the PIP-disrupting glycine mutants was obtained (**Fig. 4B**), suggesting that both decreasing and increasing the affinity to PCNA can compromise the ability of Spd1 to cause checkpoint activation.

The interaction between Spd1 and PCNA has previously been studied *in vivo* by BiFC (29, 31). In this assay, a signal is only observed when Spd1 turnover is switched *off* by inactivation of the CRL4^Cdt2^ ubiquitylation pathway. We therefore tested the ability of the PCNA-interaction mutants to generate a BiFC signal following repression of Cdt2 (**Fig 5A**). Surprisingly, none of the tested mutants, including *spd1-4G*, appeared to prevent the interaction of Spd1 with PCNA altogether. Recently, we showed that Spd1 harbours a disordered ubiquitin binding motif (DisUBM) in its C-terminus involving the residues _108_E**W**LKP**F**D_114_ (32). In fission yeast, PCNA becomes mono-ubiquitylated on K164 during unperturbed S phase, and further poly-ubiquitylated following DNA damage (55). We therefore tested the importance of these K164 modifications for Spd1 function. However, blocking PCNA ubiquitylation by introducing a *pcn1-K164R* substitution or by inactivating the Rhp18 ubiquitin protein ligase did not compromise the ability of Spd1 to cause checkpoint activation (**Fig. 5C**). On the other hand, taking out the ability of Spd1 to bind ubiquitin by mutation (*spd1^W109G^ ^F113G^* mutant (*spd1-2G3*)), significantly reduced Spd1 dependent checkpoint activation (**Fig. 5C**). Taken together, these observations suggest that ubiquitylation indeed appears to be important for Spd1-mediated checkpoint activation, although the PCNA K164 modifications seem not important for this.

**Fig. 5.**
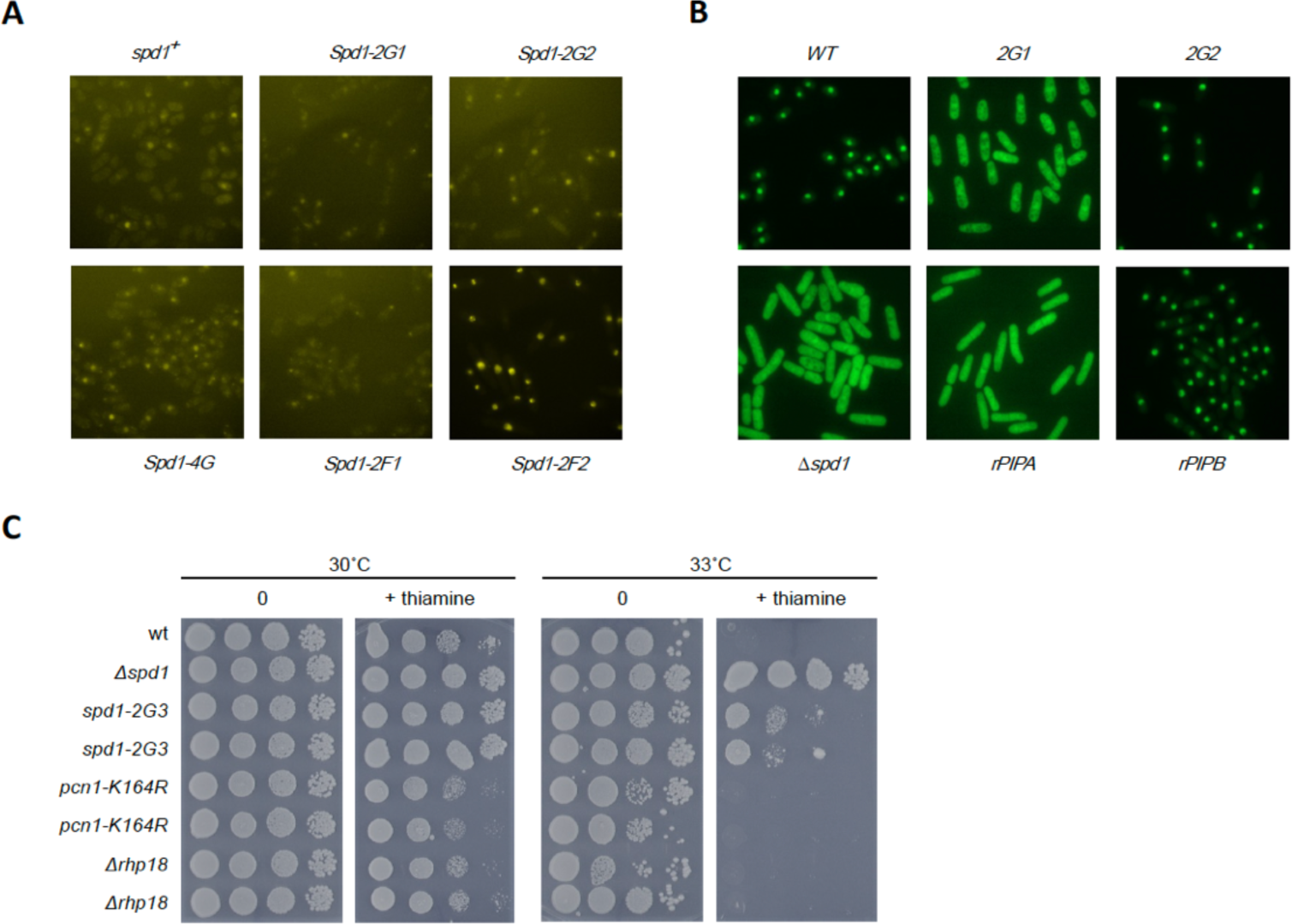
Consequences of PCNA-interaction mutants for in vivo function of Spd1. **A**) Interaction of spd1 with PCNA studied by BiFC. All strains harbour the *cdt2-TR* allele. Cells were incubated at 30°C in the presence of thiamine for three hours prior to photography. **B**) Localization of GFP-Suc22 in cells of the indicated *spd1* genotypes. **C**) Serial dilutions of strains with the indicated genotypes were spotted on plates either with or without thiamine and incubated at the indicated temperature for three days. All strains also harbour the cdt2-TR and the rad3-TS alleles.

### *In silico* analysis of full-length Spd1 binding reveals formation of compactons

The two SLiMs are separated by 14 residues and we hypothesised that they take turns in binding the same PCNA subunit, at least on a short timescale. Two other scenarios were considered: one Spd1 would bind two adjacent PCNA subunits simultaneously, or one of the PIP SLiMs would bind to one of the PCNA subunits, positioning the other SLiM close to an adjacent subunit, such that when the bound SLiM is released, the ligand jumps to the next PCNA subunit *via* a local concentration effect.

To understand and visualize the conformations of Spd1 in the PCNA bound state, Monte Carlo simulations were performed. We used the ABSINTH implicit solvent model (41) as implemented in the Campari software (42). Two different systems were simulated in which Spd1 was either bound to PCNA *via* the PIP-degron or *via* the PIP-box motif. The PCNA structure was fixed except for the eight C-terminal residues, which belong to a disordered region and somehow are involved in binding. To avoid unbinding of the disordered ligand, the Spd1 residues that interact directly with the generic ligand binding site on PCNA were also kept fixed. We added ions to neutralize the system and simulated at different NaCl concentrations.

The simulations showed a relatively compact C-terminal part of Spd1 (residues 44-124) in agreement with the *R_h_* extracted in the NMR diffusion experiments. Indeed, these 81 C-terminal residues appeared as two distinct disordered, but compact regions, which we refer to as “compactons” (**Fig. 6**). There were no secondary structure elements present or deep local energy minima, but in each compacton, a relatively large number of aromatic residues (5 and 7, respectively) formed an unstructured constantly shifting core. Mapping the distances from the terminus of Spd1 with one motif bound to site C in PCNA to the other two sites, A and B, revealed distances that were so long that they would be incompatible with rebinding of the second motif in an adjacent site (**Fig. 6B-D**). Furthermore, in the PIP-degron-bound state, the PIP-box (now constituting part of the first compacton) interacted with the interdomain connecting loop on PCNA in a dynamic manner via several different contacts in a heterogeneous (fuzzy) interaction (**Fig. 6E**) (56, 57). This weak interaction is supported by the presence of signals in the NMR spectrum of PCNA, not disappearing due to slow exchange. The residues that mainly participated in interactions with the PCNA interdomain connecting loop were from the PIP-box motif. Throughout the simulation, the N-terminal part of the protein upstream from the PIP-box, which includes the PIP-degron motif, found its way to the middle of the central channel in the clamp, where the first 16 N-terminal residues formed a small compacton without any direct interactions with PCNA (**Fig. 6E**). This could explain the effect on the chemical shift changes in the N-terminal part of Spd1.

**Fig. 6.**
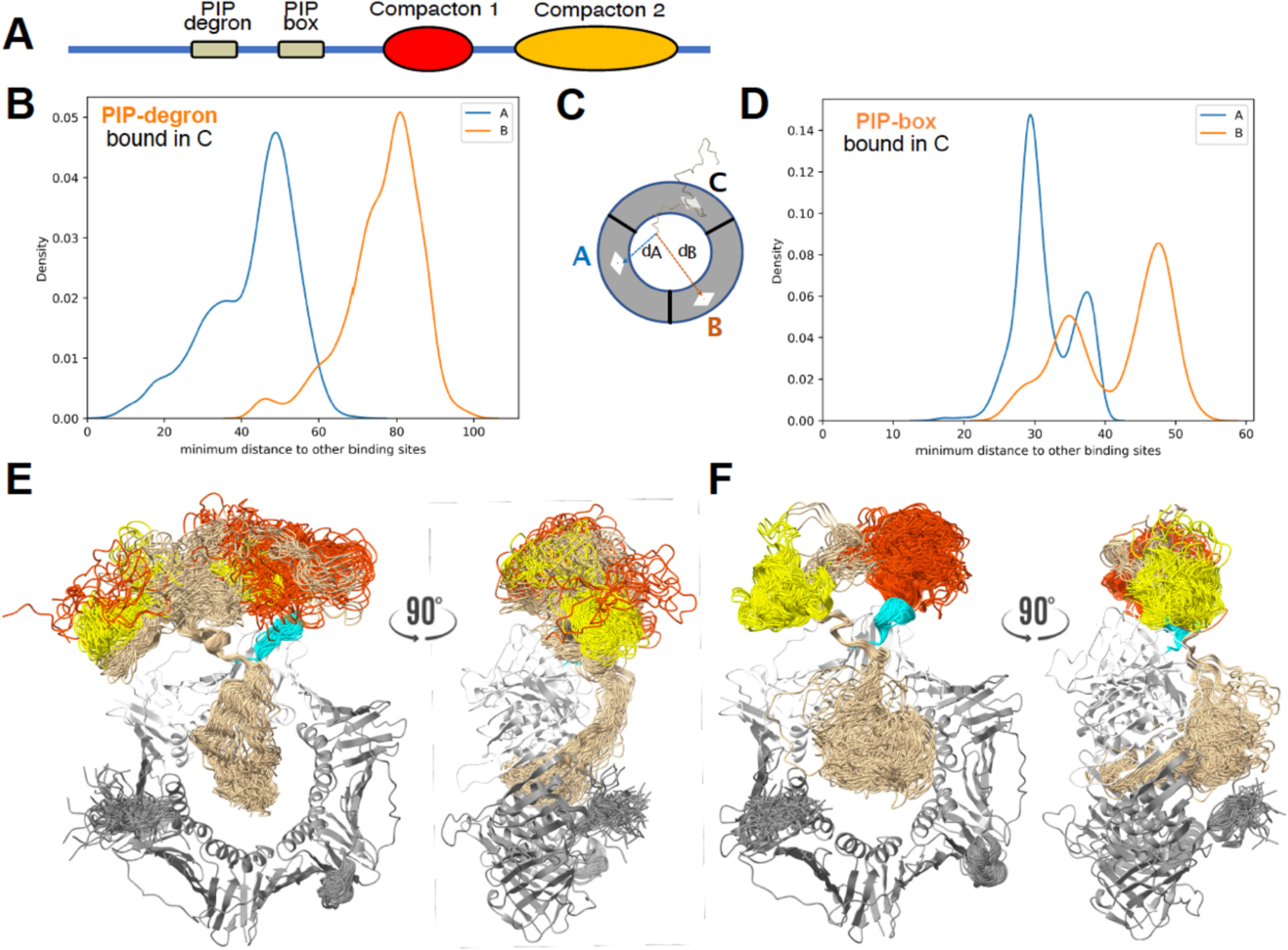
*In silico* analysis of Spd1 binding to PCNA reveals compacton formation and different ensembles depending on the active SLiM. **A)** Schematic overview of the fully disordered, 124 amino acid residues Spd1 protein. **B**). The probability of finding the PIP-box SLiM at different distances from the PIP binding pocket of subunits A (blue line) or B (orange line). **C**) Schematic representation of the principle plotted in B) and D). **D**) The probability of finding the PIP-degron as a function of distance to the PIP binding pockets on subunits A (blue) and B (orange). **E**) 100 trajectories of the simulation of the PIP-degron bound state superposed. The C-terminal disordered tail from the C-subunit of PCNA interacts with the compactons, and is coloured light blue. The compactons are red and yellow according to A). **F**) Like E, only with Spd1 bound to PCNA via the PIP-box SLiM. The C-terminal compactons (red and yellow according to A)) are more distinct, the PCNA C-terminal tail interacts exclusively with compacton 1. The N-terminal Spd1 chain cover the central opening in the PCNA toroid.

In the PIP-box bound state, the PIP-degron was also found inside the channel along with the rest of the N-terminus. (**Fig. 6F**) In this case there were interactions with the positively charged inner brim, especially via residues from the PIP-degron SLiM. The first C-terminal compacton was disrupted because part of it, the PIP-box, was now bound to the hydrophobic binding pocket in PCNA. The second compacton remained undisturbed during the simulation. We emphasize that the simulations did not include DNA.

The computer simulated structures of both trajectories do not exclude allovalency, but of the three scenarios mentioned, it seems probable that the SLiMs take turn in binding to the same subunit. However, the two complexes differ so much in the entire Spd1 context that it is tempting to speculate two different biological roles for the two states rather than providing avidity; biological roles that may depend on the presence of other factors, *e.g.,* but not restricted to, Suc22^R2^.

## Discussion

In the present study, we have characterized the interaction between Spd1 and PCNA in detail. In addition to the formerly identified PIP-degron (29), we report the existence of a second PIP-motif – a PIP-box - in Spd1, and we show that these two tandemly positioned low affinity sites exchange rapidly on PCNA. When we reduced the affinity of either or both sites to PCNA, we compromised the ability of Spd1 to cause checkpoint activation i*n vivo*, consistent with the interaction with PCNA indeed being important for Spd1 function. In an effort to restore the checkpoint response, we designed and confirmed higher affinity PCNA binding by introducing the two canonical aromatic amino acids to both Spd1 PIP-SLiMs, one by one. Much to our surprise, these mutations resulted in the same checkpoint-deficient phenotype as the glycine mutations that abolish binding to PCNA. This result is inconsistent with the PCNA brake model, as one would expect increased checkpoint activation with enhanced binding. Moreover, the *K_d_* for p21 binding to PCNA is at least three orders of magnitude lower than that of Spd1(8), suggesting that Spd1 simply binds PCNA too weakly to block replication.

How then does Spd1 binding to PCNA enable checkpoint activation? The fact that both an increase and a decrease in the affinity between the two compromises checkpoint activation suggests that there exists a fine-tuned balance of affinities and molar concentrations of Spd1 and its ligands, PCNA and Suc22^R2^. Indeed, both sites are conserved, both in the homologues Spd2 and in other schizosaccharomyces species (24). Disturbing that balance by tampering with optimized coupled affinities hinders proper checkpoint activation. Either increasing or decreasing the affinity between Spd1 and PCNA, or overexpressing Suc22^R2^ can accomplish this (58).

When studying variants incapable of binding to PCNA with either one or both SLiMs compromised, we found that both sites are important for function, but that the double glycine mutant in the PIP-degron motif had a more severe effect (data not shown). Importantly, although the Suc22^R2^ binding site on Spd1 is not known in molecular details, triple-alanine mutations along the entire sequence of Spd1 pinpointed it to the HUG-domain region comprising residues Q30-Y63 (25), a region also encompassing the PIP-degron. We therefore propose that the slightly different biological responses to knocking out either the PIP-degron or the PIP-box by introducing two glycine residues in either SLiM, are caused by the overlap of the binding sites for Suc22^R2^ and PCNA. The PIP-degron mutant probably not only loses its ability to bind PCNA, but also compromises interaction with Suc22^R2^. This is supported by our observation that the *spd1-2G1* and *spd1-2F1* mutants both abolish Spd1-mediated nuclear localization of Suc22^R2^. Further, we note that the insertion of the glycine in the PIP-degron (G37), disrupts its canonicity, but is important for RNR binding, as, when mutated, it results in loss of ability to restrain RNR activity (25). Thus, we speculate this glycine to be key to RNR interaction, compromising the canonicity of the PIP-degron.

Monte Carlo simulations of the two PCNA-Spd1 complexes, bound either via the PIP-box or via the PIP-degron, revealed two very different structural ensembles. In the PIP-degron complex, the PIP-box SLiM is close to the PCNA binding pocket, whereas the PIP-degron is far from the binding pocket when the PIP-box is bound. This is in accord with observations published by Salguero *et al*. (29), who discovered the PIP-degron, but not the PIP-box. They found that Spd1-variants missing the PIP-degron but retaining the PIP-box, underwent only very modest degradation, whereas variants missing the box but having the degron were degraded efficiently.

Why then are two PIP motifs needed in Spd1 and why the weak affinities? Of the tandemly positioned SLIMs, the composite SLiM involving both the PIP-degron and the so-far undefined Suc22^R2^ binding motif may have compromised PCNA binding to an extent that has led to the evolution of a second SLiM, which jointly achieve the required affinity. They are separated by 13 residues, which made us hypothesise an avidity effect, meaning that even if the affinity is low for either site, the proximity raises the effective concentration (54), increasing the apparent affinity. This is observed to some extent. The affinity for the individual PIP SLiMs are low (both around 3 mM), whereas the affinity for the full-length Spd1 protein is ∼300 µM, a 10-fold decrease in *K_d_*. Since the flanking regions have been shown to participate in disordered binding to PCNA (8), some of the effect must be attributed to this. On this background, the avidity effect seems modest. However, both Spd1 and PCNA may be targets for post-translational modifications like phosphorylation and ubiquitylations, as described for PCNA (59, 60). Using recombinant proteins as in the NMR experiments presented here, does not capture affinity tuning by posttranslational modifications. The fact that the *spd1-4G* mutant still show some degree of interaction with PCNA in our BiFC assay, suggests that *in vivo* modifications also contribute to the interaction. Our observation that the *spd1^W109G^ ^F113G^* mutant – defective in ubiquitin binding – is partially defective in checkpoint activation suggests that protein-ubiquitylation somehow plays a role in the process.

The observations reported here make it tempting to speculate that the two sites have different biological roles, as in the case of human Pol η mentioned in the introduction. Recently, structures of the PCNA-bound Pol δ were published, revealing the presence of two PIP-boxes called iPIP and cPIP (61). cPIP is capable of binding PCNA as well as a primase. In the active complex, iPIP binds PCNA and cPIP binds the primase. Thus, it is possible that the PIP-box of Spd1 may bind a yet unknown target. One of the observations made in the simulations, and by NMR, is the low solubility and the presence of compact states of Spd1 and the presence of compactons involving many aromatic residues, two of which are part of the PIP-box. The unusual large-scale self-interaction may be related to a property of Spd1 to form higher order structures as seen for many multivalent proteins and IDPs in the formation of condensates, leading to phase separation (62, 63). This has not yet been reported for Spd1, or for any of the orthologs in *S. cerevisiae* (Sml1, Dif1, Hug1), but one may speculate that such higher order dynamic structures will be of relevance to the regulation of both PCNA and Suc22^R2^ and for activating the checkpoint. If this is the case, then increasing the number of aromatic residues in Spd1 as with the restored canonical PIP-variants (Spd1-2F1 and Spd1-2F2) may lower the saturation concentration to form condensates. This could be a clue to the reason for the absence of the canonical aromatic residues and may also explain the low affinity, which will increase in the high concentrations within the condensates. These speculations remain to be addressed.

The observations presented here point to a delicate system competing for ligands at the hub; a game of affinities and a balance of concentrations, but also suggesting that binding kinetics (*k_off_*/*k_on_*), post translational modifications and context contribute substantially. PCNA binds numerous factors from polymerases to DNA methyl transferases. The exchange of these factors depends on high specificity and relatively low affinity, as in the case of Spd1, but also on availability (64). This ensures the establishment of a system, which at the same time is hyper complex, dynamic and precise, but highly prone to fast regulation. We suggest that low-affinity PCNA binding ligands with composite PCNA-binding SLiMs are relevant in many other PCNA-binding ligands that so far have gone under the PIP-box radar, because they are invisible to current sequence analysis.

## Data availability

The assignments of *S. pombe* PCNA have been deposited in the BMRB data bank under the accession code 51376. The coordinates and structure factors for the crystal structure of the complex of S. pombe PCNA and Spd1^27-46^ are deposited in the PDB under the accession code 6QH1. All input files for the MC simulations can be downloaded from https://github.com/rcrehuet/spd1_PCNA

## Acknowledgments

We thank Signe A. Sjørup for excellent technical assistance and the many students struggling in the early phases of Spd1 handling.

## Open Access

This article is distributed under the terms of the Creative Commons Attribution 4.0 International License (http://creatIvecommons.org/licenses/by/4.0/), which permits unrestricted use, distribution and reproduction in any medium, provided you give appropriate credit to the original author(s) and the source, provide a link to the Creative Commons license, and indicate if changes were made.

## SUPPLEMENTAL INFORMATION

**Supplementary Fig. S1:**
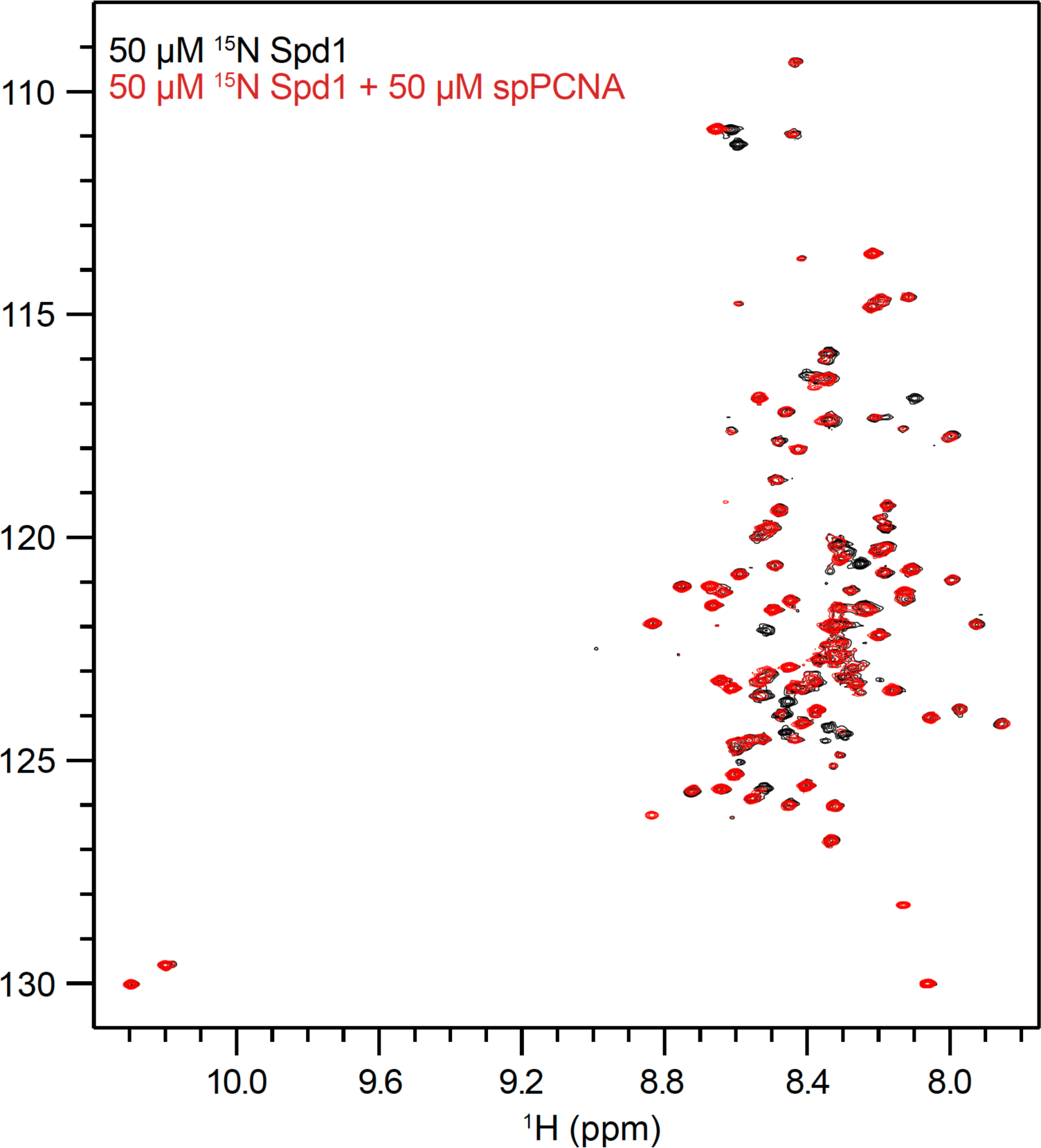
^1^H,^15^N-HSQC spectra of ^15^N-Spd1 with and without addition of spPCNA.

**Supplementary Fig. S2:**
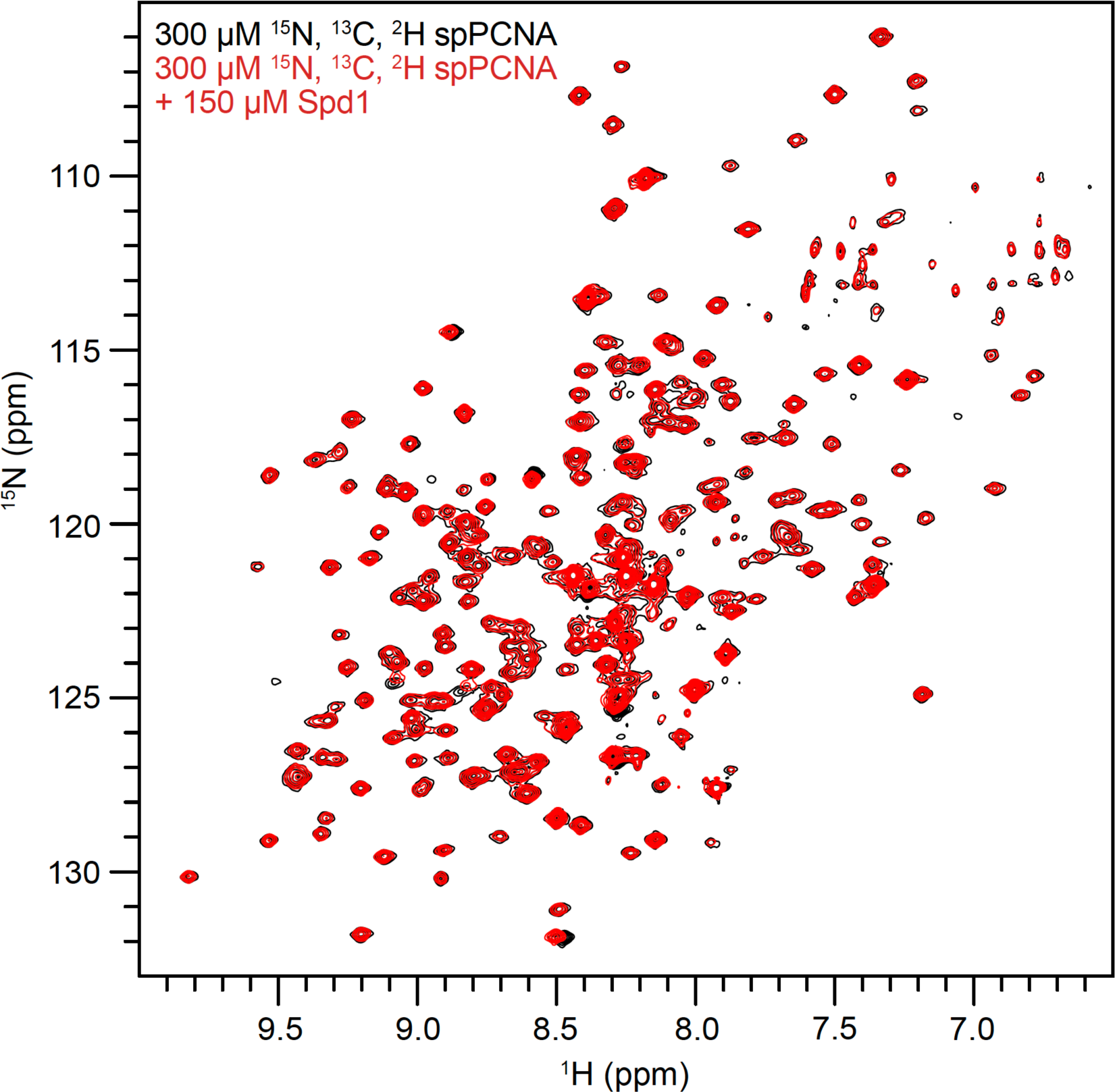
^1^H,^15^N-HSQC spectra of ^15^N,^13^C,^2^H-spPCNA with and without addition of Spd1.

**Table S1.**
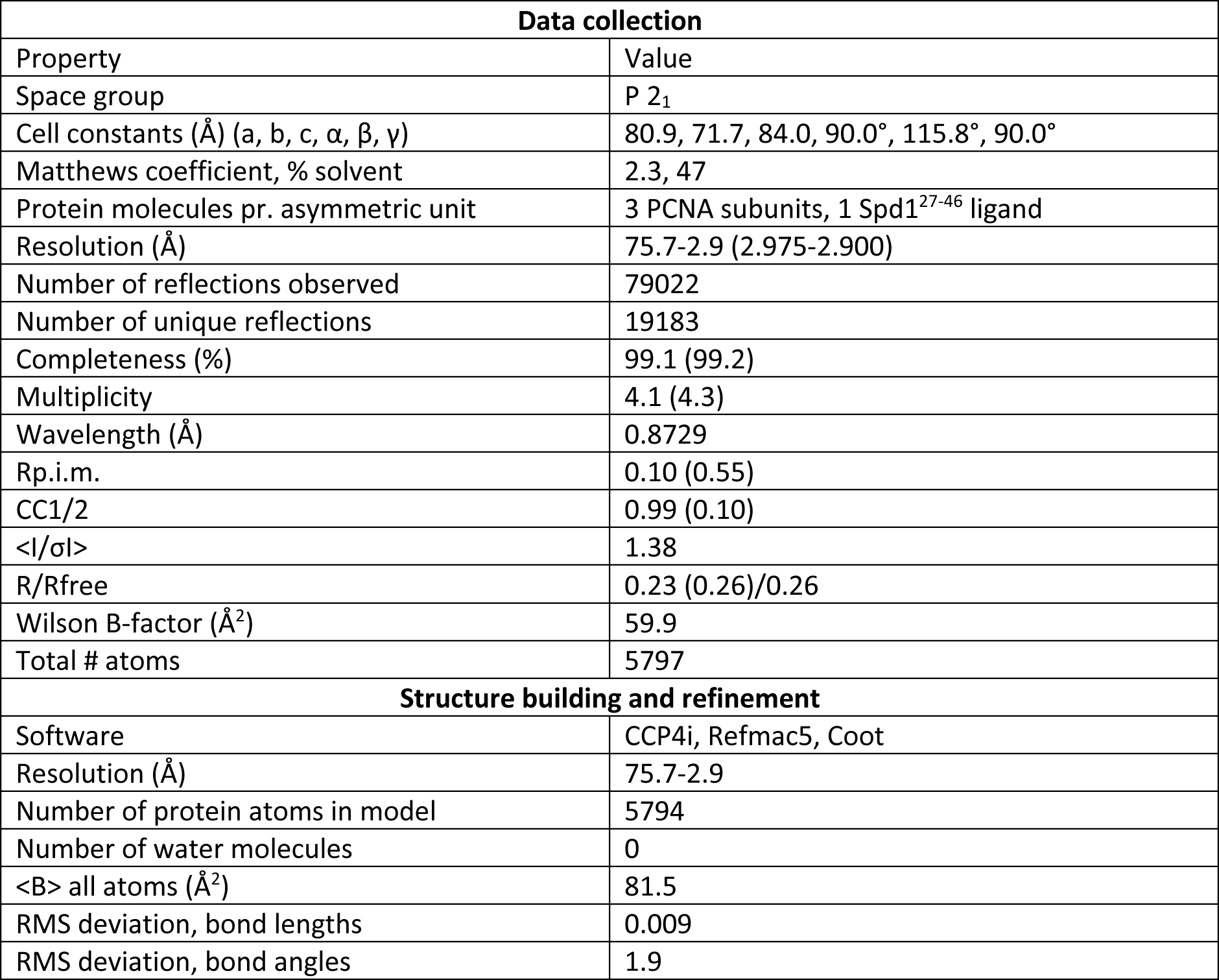
Data collection and refinement data for PCNA:Spd1. Values in parenthesis refer to data from the outer resolution shell (3.06-2.90 Å).

**Table S2.** Fission yeast strains.

| Strain: | Genotype: |
| --- | --- |
| Eg3439 | *h<sup>+</sup> leu1+::nmt41-cdt2 cdt2::ura4+ ura4-D18 Δspd1::hygro rad3-ts* |
| Eg3490 | *h<sup>+</sup> leu1+::nmt41-cdt2 cdt2::ura4+ ura4-D18 rad3-ts* |
| Eg4191 | *h<sup>+</sup> leu1+::nmt41-cdt2 cdt2::ura4+ ura4-D18 rad3-TS spd1-L33G V36G* |
| Eg4193 | *h<sup>+</sup> leu1+::nmt41-cdt2 cdt2::ura4+ ura4-D18 rad3-TS spd1-F55G Y58G* |
| Eg4549 | *h<sup>+</sup> leu1+::nmt41-cdt2 cdt2::ura4+ ura4-D18 spd1-V36F G37F rad3-ts* |
| Eg4553 | *h<sup>+</sup> leu1+::nmt41-cdt2 cdt2::ura4+ ura4-D18 spd1-F55L N59F rad3-ts* |
| Eg4035 | *h<sup>-</sup> GFP-suc22* |
| Eg4031 | *h<sup>-</sup> GFP-suc22 Δspd1::hygro* |
| Eg4892 | *h<sup>+</sup> spd1-L33G V36G GFP-suc22* |
| Eg4894 | *h<sup>+</sup> spd1-F55G Y58G GFP-suc22* |
| Eg4900 | *h<sup>+</sup> spd1-V36F G37F GFP-suc22* |
| Eg4904 | *h<sup>+</sup> spd1-F55L N59F GFP-suc22* |
| Eg3947 | *h<sup>+</sup> leu1+::nmt41-cdt2 Δcdt2::ura4+ ura4-D18 VN173-pcn1::kmx spd1-VC155::nat* |
| Eg4926 | *h<sup>+</sup> leu1+::nmt41-cdt2 cdt2::ura4+ ura4-D18 VN173-pcn1::kmx spd1-L33G V36G-VC155::nat* |
| Eg4927 | *h<sup>+</sup> leu1+::nmt41-cdt2 cdt2::ura4+ ura4-D18 VN173-pcn1::kmx spd1-F55G Y58G-VC155::nat* |
| Eg4928 | *h<sup>+</sup> leu1+::nmt41-cdt2 cdt2::ura4+ ura4-D18 VN173-pcn1::kmx spd1-L33G V36G-F55G Y58G-VC155::nat* |
| Eg4929 | *h<sup>+</sup> leu1+::nmt41-cdt2 cdt2::ura4+ ura4-D18 VN173-pcn1::kmx spd1-V36F G37F-VC155::nat* |
| Eg4930 | *h<sup>+</sup> leu1+::nmt41-cdt2 cdt2::ura4+ ura4-D18 VN173-pcn1::kmx spd1-F55L N59F-VC155::nat* |
| Eg4514/<br>Eg4515 | *h<sup>+</sup> leu1+::nmt41-cdt2 cdt2::ura4+ ura4-D18 VN173-pcn1::kmx spd1-W109G F113G-VC155::nat* |

